# Image-based temporal profiling of autophagy-related phenotypes

**DOI:** 10.1101/2023.03.27.534404

**Authors:** Nitin Sai Beesabathuni, Eshan Thilakaratne, Priya S. Shah

## Abstract

Autophagy is a dynamic process that is critical in maintaining cellular homeostasis. Dysregulation of autophagy is linked to many diseases and is emerging as a promising therapeutic target. High-throughput methods to characterize autophagy are essential for accelerating drug discovery and characterizing mechanisms of action. In this study, we developed a highly scalable image-based profiling approach to characterize ∼900 morphological features at a single cell level with high temporal resolution. We differentiated drug treatments based on morphological profiles using a random forest classifier with ∼90% accuracy and identified the key features that govern the classification. Additionally, temporal morphological profiles accurately predicted biologically relevant changes in autophagy after perturbation, such as total cargo degradation. Therefore, this study acts as proof-of-principle for using image-based profiling to differentiate autophagy perturbations in a high-throughput manner and identify biologically relevant autophagy phenotypes, which can accelerate drug discovery.

## Introduction

Macroautophagy (hereafter referred to as autophagy) is an intracellular process that recycles damaged and unwanted cellular components to maintain homeostasis. In autophagy, specialized double-membraned structures called phagophores grow to surround cellular contents such as proteins and organelles to form autophagosomes. Following the fusion of autophagosomes with lysosomes, which forms autolysosomes, the cellular cargo is degraded by lysosomal enzymes. This recycling process results in the generation of free amino acids, fatty acids, and energy for biosynthetic pathways, and can help turn over damaged proteins and organelles [1].

Autophagy is linked to many diseases and disorders [2–4]. For example, failure to clear damaged proteins and organelles from long-lived cells such as neurons results in cytotoxicity and neurodegeneration [5]. Upregulating autophagy can help cancer cells survive in nutrient-depleted environments [6]. Given its role in disease, fine-tuning autophagy to reverse or prevent disease is of great interest [7,8]. Quantitative and holistic characterization of autophagic activity is essential to this effort. Microtubule-associated protein 1A/1B-light chain 3 (LC3) is the most commonly used marker to monitor autophagy [9]. LC3 is often tagged with fluorescent molecules to enable quantitative measurements of autophagy phenotypes[10]. Nevertheless, quantitative measurements of autophagy at a single cell level are primarily limited to autophagy vesicle count using fluorescence microscopy and image cytometry, or mean intensity of the cell using flow cytometry [10]. Although these measurements have enabled high-throughput, quantitative, and sensitive detection of autophagy, they fail to capture the autophagy state comprehensively. The two main reasons are 1) vesicle count and mean intensity alone do not inform the rate of flow of cargo through the pathway, often referred to as autophagy flux [10], and 2) other properties such as shape, size, and distribution of autophagy vesicles over time, as well as whole-cell morphological changes, are often not analyzed, leaving vast amounts of phenotypes uncharacterized.

We recently developed a quantitative framework to measure autophagy rates to address the first challenge [11]. Our approach expands on the quantitative steady-state autophagy flux analysis originally developed by Loos and colleagues [12]. Our instantaneous rate approach allows us to measure the rate of each step in the pathway under dynamic conditions. By uncoupling measurements of the rate of autophagosome formation from the rate of autophagosome-lysosome fusion and autolysosome turnover, we can better resolve the direct mechanisms of action and long-term feedback loops in response to environmental perturbations (*e*.*g*., chemicals or nutrients). We can also resolve the total degradative capacity of dynamic and adaptive systems, which can assist in precisely fine-tuning autophagy flux. However, this approach involves destructive sampling, limiting the scalability of this method to characterize thousands of conditions in a high-throughput manner.

Image-based profiling facilitates the simultaneous quantification of various morphological features of sub-cellular components at a single cell level. Characterization of autophagy-related phenotypes using image-based profiling approaches has been performed for various applications such as the characterization of small molecule regulators and genetic modulators [13–15]. These studies highlight the potential of using image-based profiling for high-throughput autophagy characterization. However, these studies were performed at a single time point after fixing cells, limiting our understanding of dynamic changes in autophagy phenotypes.

To address these challenges, we created an image analysis pipeline to systematically characterize temporal changes in autophagy-related morphological phenotypes at a single cell level. We investigated changes in morphological features following the addition of two common, well-characterized small molecule autophagy modulators rapamycin and wortmannin. We examined the key morphological features differentially modulated under various treatments as a function of time. Using a random forest classifier, we identified the features with high importance that can be used to differentiate these treatments. We also used a profile similarity approach to test the possibility of characterizing autophagy regulation based on morphological features. Compared to canonical measurements of autophagy vesicle dynamics, the inclusion of additional morphological features captured autophagy modulation more efficiently. Overall, this study serves as proof-of-concept for using image-based profiling to characterize autophagy. This approach has the potential to map novel drugs to known mechanisms of action and facilitate high-throughput drug characterization by linking complex morphological responses to biologically relevant cellular outputs.

## Results

### Experimental setup and image-based profiling pipeline

We quantified various morphological features using the pHluorin-mKate2-LC3 system[16] by reanalyzing images that were previously collected to measure autophagy rates [11] [**Fig 1A**]. An illustrative image of the change in cellular morphology is shown in **Fig 1B**, where wortmannin treatment caused void-like regions in the cells, while rapamycin caused a decrease in the signal intensity of cells. We developed an image analysis pipeline to segment and track individual cells over time [**Fig 1C-D**]. Along with autophagosome and autolysosome numbers, three main morphological properties (structure, intensity, and texture) were quantified at a single cell level for whole cell (Cell), autophagosomes (AP), and autolysosomes (AL) [**Fig 1E**]. Approximately 900 features for each cell were quantified for the entire time course of the experiment. The detailed pipeline is shown in **Figure S1A** and described further in the methods.

**Figure 1:**
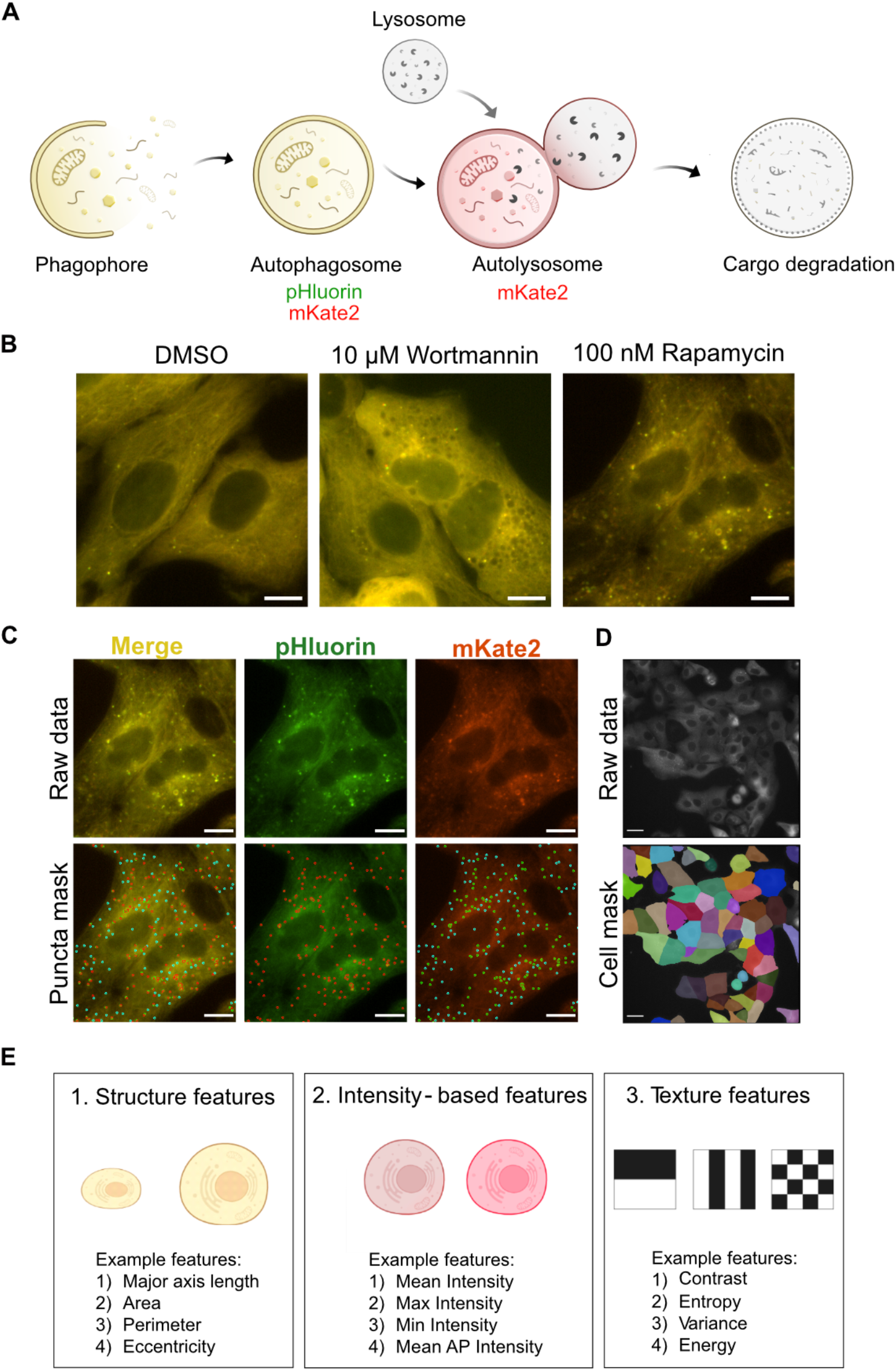
Experimental and image analysis pipeline to quantify autophagy-related phenotypes. **A:** pHluorin-mKate2-LC3 was used for tracking autophagosomes and autolysosomes. **B:** Representative images of change in morphology of cells after treatment with 100 nM rapamycin and 10 μM Wortmannin for 6 hours. The scale bar represents 10 μm. **C:** Representative images of spot detection tool for detecting autophagosomes (red spots in the pHluorin channel) and autolysosomes (cyan spots in the mKate2 channel). Scale bar represents 10 μm. **D:** Representative image of segmented cell mask used for tracking individual cells. **E:** Three main categories of morphological features were extracted at a cellular level as well as an autophagy vesicle level.

### Temporal profiling of morphological features during autophagy induction and inhibition

We characterized morphological changes in cells treated with high concentrations of rapamycin and wortmannin. We previously observed that rapamycin and wortmannin treatment increased and decreased overall autophagy flux, respectively [11]. We first identified the features that varied significantly compared to DMSO-treated cells at each time point for both treatments. All features with a median modified Z score ≥ 0.5 with an adjusted P-value < 0.05 were considered significant. A representative analysis is shown at 0.5 hours after treatment with 10 μM wortmannin and 100 nM rapamycin [**Fig 2A-B**]. Identical analysis was performed on the same cells before treatment to confirm that a threshold of 0.5 removes any false positive features [**Fig S1B-C**]. We compared measurements from our image analysis pipeline to those obtained from flow cytometry. Data from both methods showed a significant decrease in mean fluorescence intensity following treatment with rapamycin for 6 hours [**Fig S1D-E**], confirming the consistency of the imaging pipeline with other methods commonly used to screen autophagy responses at a single cell level.

**Figure 2:**
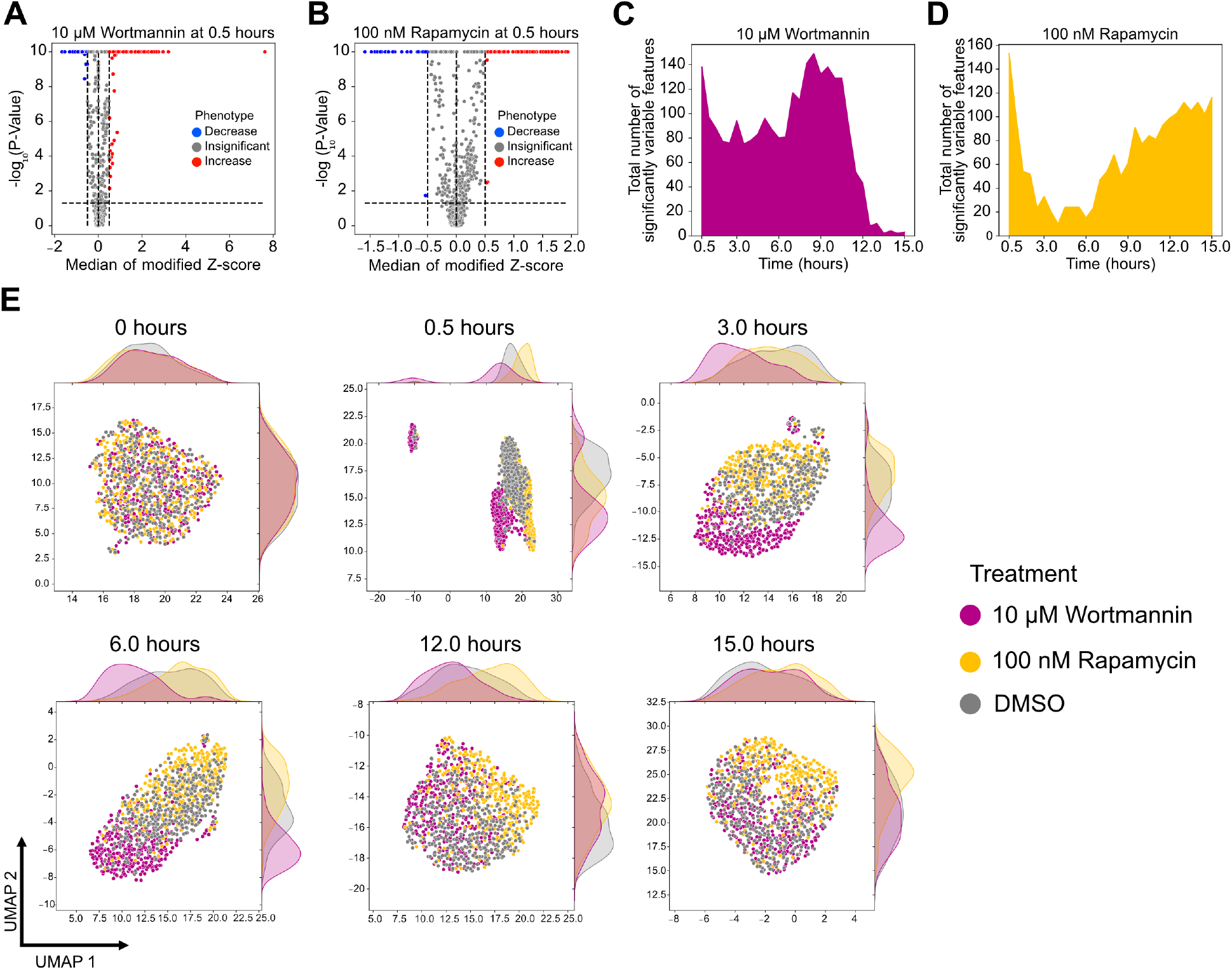
Temporal change in morphological features after rapamycin and wortmannin treatment. **A-B:** Volcano plots of cellular features after 30 minutes of treatment with **A:** 10 μM wortmannin and 100 nM rapamycin, respectively. **C-D:** Cellular features that varied significantly as a function of time for 10 μM wortmannin and 100 nM rapamycin, respectively **E:** UMAP of cells treated with DMSO, 100 nM rapamycin, and 10 μM Wortmannin at different time points. A minimum of 300 cells were analyzed for each condition. Features that varied significantly with a median modified Z-score ≥ 0.5 at specific time points were used for generating the UMAPs. For 0 hours, features from 0.5 hours were used.

We next analyzed the number of features that varied significantly with time [**Fig 2C-D**]. Wortmannin and rapamycin treatment both led to dramatic increases in variable features immediately following treatment. Wortmannin-treated cells maintained this increase until ∼12 hours. Conversely, rapamycin-treated cells showed a rapid decrease following initial peak and then a gradual increase in variable features that was maintained even at 15 hours. We also analyzed a lower concentration of wortmannin and wortmannin/rapamycin combination treatments using a similar approach [**Fig S2A-C**] and observed a concentration dependent effect of wortmannin on cellular morphology dynamics. We visualized the single cell landscape for DMSO, wortmannin, and rapamycin treatment at multiple time points using uniform manifold approximation and projection (UMAP) [17] [**Fig 2E**]. While cells from different treatments were distributed evenly throughout a single cluster before treatment, they formed distinct clusters based on treatment in as little as 30 minutes. Wortmannin-treated cells were more easily separated from DMSO-treated cells at all time points before 12 hours compared to rapamycin. After 12 hours, the wortmannin-treated cells clustered together with DMSO-treated cells which is consistent with the decrease in the differential features at later time points [**Fig 2C**]. Conversely, rapamycin-treated cells re-segregated from DMSO-treated cells beginning at 12 hours and maintained this segregation for the duration of the experiment. In conclusion, dynamic changes in morphological features can be used to visualize distinct cellular responses in an unsupervised manner.

### Feature importance in distinguishing rapamycin and wortmannin treatment using a random forest classifier

Identifying governing features that distinguish one drug from another in the autophagy response would be a powerful tool for dissecting mechanism of action and discovery of novel phenotypes. Toward this end, we constructed a random forest model to identify the primary feature set that could be used to differentiate rapamycin and wortmannin treatments. The model was trained using all variable features after removing correlated features (see Material and Methods). The model achieved an F_1_ score of 0.89 ± 0.033 [**Fig 3A**]. This random forest classifier was then examined using Shapely Additive Explanations (SHAP) values to establish the importance of each feature in differentiating the treatments [18]. A higher absolute SHAP value represents a higher importance in classifying the treatments. We grouped the features based on biological category (Cell, AP, and AL) and morphological property (structure, intensity, and texture), then calculated the cumulative as well as average importance for each group in classifying wortmannin and rapamycin treatments [**Fig 3B-C**]. Cumulative importance represents the sum of absolute SHAP values of all features in each group while average importance represents the mean of absolute SHAP values of all features in each group. Autophagosome texture and structure groups had a high cumulative importance for both treatments but contained a large fraction of the variable features which might contribute to the overall cumulative value. Vesicle numbers and Cell/Intensity groups had a low number of features but contributed substantially towards the classification of both treatments indicated by the high average importance value. We analyzed individual features that influence the classification, with the top 15 shown in **Figure 3D**. The mean intensity of the cell (mean_intensity) at 0.5 hours and autophagosome texture (contrast_mean) were key features in differentiating wortmannin from basal conditions. Initial autophagosome number (AP_number) and autophagosome structural features such as maximum autophagosome area (max_AP_area) and maximum autophagosome Zernike moment (max_AP_Zernike moment_10) were important in classifying rapamycin treatment.

**Figure 3:**
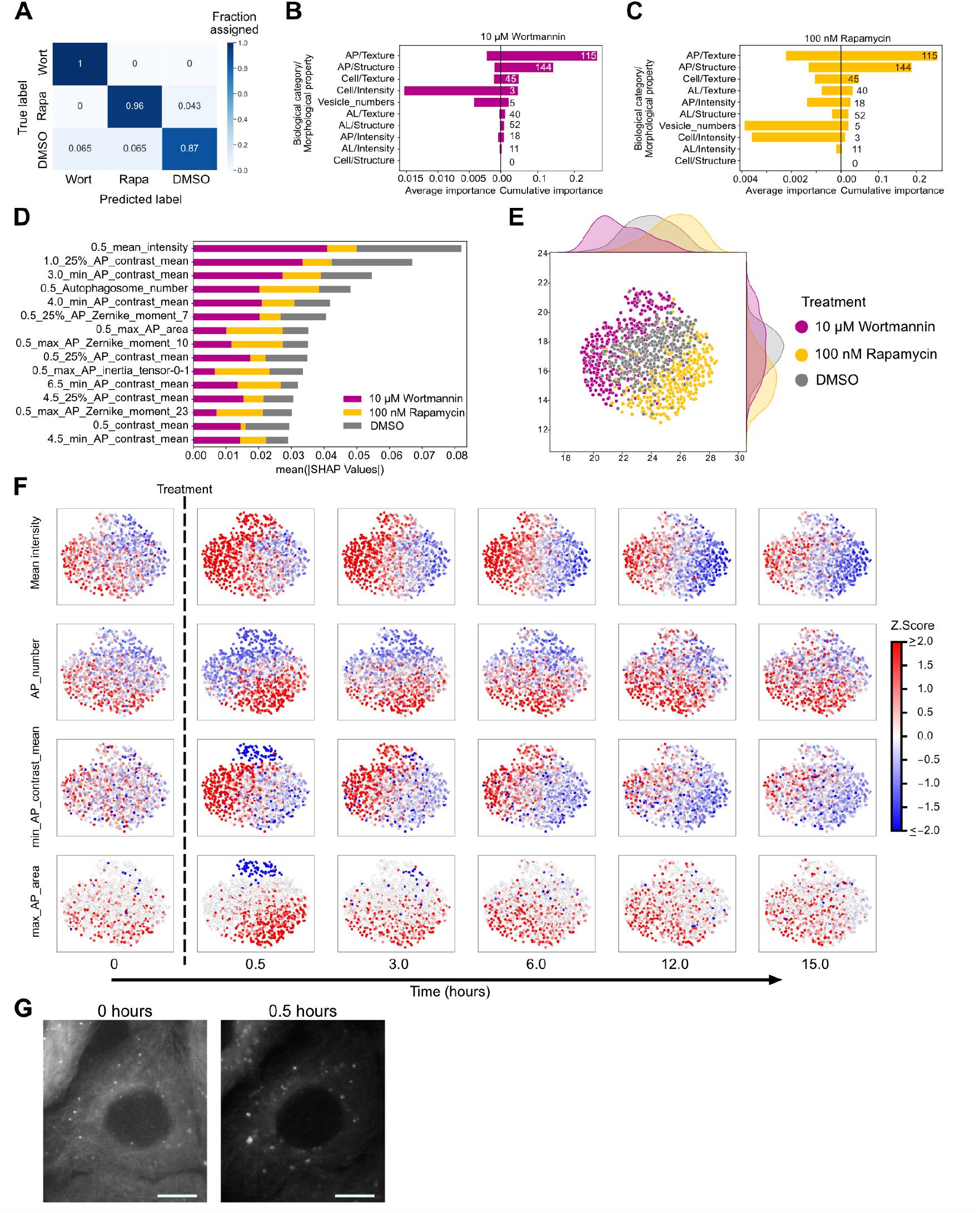
Differentiating rapamycin and wortmannin treatments and governing features. **A:** A confusion matrix to visualize the performance of the random forest model in classifying 10 μM wortmannin (Wort) and 100 nM rapamycin (Rapa) and DMSO treatments. A minimum of 70 cells for each condition were used for testing the accuracy of the model. **B-C:** Cumulative and average important of feature categories in classifying wort and rapa treatments, respectively. **D**: The top 15 features with the highest mean absolute SHAP values. **E**: UMAP of individual cells constructed using variable features from all time points. A minimum of 300 cells were analyzed for each condition. **F**: Change in feature values at a single cell level with time after treatment with rapa and wort. **G**: Representative image of increase in autophagosome area after treating with rapa for 30 minutes. Green channel (pHluorin) images are shown in grayscale. Scale bar represents 10 μm.

We generated an UMAP of the dynamic cellular landscape to examine the impact of rapamycin and wortmannin on high importance features over time [**Fig 3E**]. The plot shows a clear separation between cells subjected to the two treatments, suggesting different mechanisms of action. We visualized mean_intensity, AP_number, minimum value of autophagosome contrast mean per cell (minimum_AP_contrast_mean), and max_AP_area to understand changes based on treatment and time. Wortmannin and rapamycin had opposing effects on initial mean_intensity and AP_number, consistent with their impacts on autophagy rates [11,19]. min_AP_contrast_mean, a strong indicator of wortmannin treatment, increased for a major fraction of wortmannin-treated cells. max_AP_area increased after rapamycin treatment, indicating that bigger autophagosomes are formed after rapamycin treatment [**Fig 3G**]. This analysis uncovered novel phenotypes and features that can be used for classifying treatments based on morphological properties.

### Temporal morphological profiling accurately predicts the dynamic change in autophagy modulation

We subsequently assessed the predictive value of temporal morphological profiling for characterizing autophagy perturbation without directly measuring autophagy rates using lysosomal inhibitors. We tested 6 conditions − 10 μM Wortmannin, 1 μM Wortmannin, 100 nM rapamycin, 10 μM Wortmannin with rapamycin, and 1 μM Wortmannin with rapamycin. These conditions previously provided a range of autophagy perturbation over 15 hours [11]. We used image-based profile matching through profile correlation [**Fig 4A**] to assess performance because this was recently demonstrated to be a promising approach for characterizing perturbations [20,21]. Image-based profile correlation was then compared to previously quantified biologically relevant changes in autophagy. Overall cargo degraded was used as the standard because it is a single measurement that captures the cumulative change in autophagy over time [**Fig 4A**] [11].

**Figure 4:**
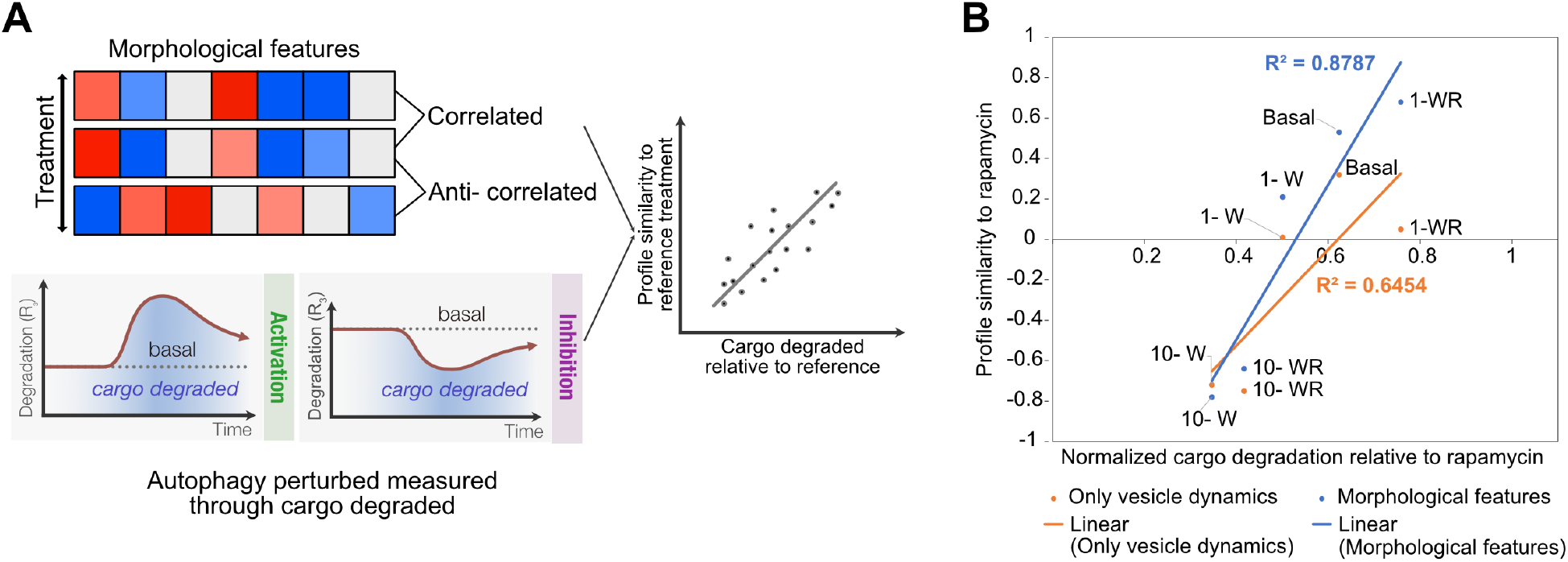
Image-based profile similarity accurately captures autophagy modulation. **A:** Cartoon illustration comparing profile similarity to cargo degradation. **B:** Correlation between profile similarity and normalized cargo degradation. Abbreviations: 10 μM Wortmannin (10-W), 1 μM Wortmannin (1-W), 100 nM rapamycin (Rapa), 10 μM Wortmannin with rapamycin (10-WR), and 1 μM Wortmannin with rapamycin (1-WR).

We compared the performance of high-dimensional image-based profiling to the less complex but more intuitive measurement of autophagy vesicle dynamics, which is commonly analyzed in autophagy studies [**Fig S3**]. Relationships between treatments identified using hierarchical clustering on morphological features were distinct to those identified based solely on autophagy vesicle dynamics [**Fig S4A-B**]. We then used profile correlation to quantitatively assess the accuracy of using either of these measurements in capturing biologically relevant changes in autophagy [**Fig S4C-D**]. Profile similarity correlations and normalized cargo degradation were calculated relative to rapamycin treatment to provide a maximum linear dynamic range with a well characterized autophagy modulator for both measurements [**Fig S4C-E**]. We observed a better linear fit between normalized cargo degradation and profile similarity measured using a comprehensive set of morphological features (R^2^= 0.8787) compared to profile similarity measured using just autophagy vesicle dynamics (R^2^= 0.6454) [**Fig 4B**]. Although simple, this analysis shows the potential of using image-based temporal profiling, which includes the measurement of non-intuitive morphological features, to comprehensively evaluate autophagy perturbation and relate this to biologically relevant cellular outputs.

## Discussion

Measuring autophagy is challenging due to its dynamic and multistep nature. Established methods accurately measure each autophagy step in the form of rates with high sensitivity, but rely on destructive sampling by the addition of lysosomal inhibitors and limit scalability [11,12]. High-throughput characterization of autophagy would enable accelerated drug screening and a better understanding of the underlying mechanisms. Even so, existing high-throughput methods involve destructive sampling and limits the ability to associate a phenotype with the temporal change in the autophagy state at the single cell level [13,22]. For example, the governing features of autophagy-dependent cell death are not fully understood. Monitoring various autophagy parameters, such as the change of different rates over time, to derive a correlation between autophagy and cell death cannot be achieved using existing methods since the same cell cannot be followed to the point of cell death using these approaches. Image-based morphological profiling offers a comprehensive and high-throughput approach for characterizing cellular phenotypes and perturbations with single cell trajectories [23,24]. We performed a proof-of-concept study to evaluate the potential of using image-based profiling for characterizing autophagy modulation.

We developed a random forest classifier with an accuracy of ∼90% to differentiate rapamycin, wortmannin and DMSO conditions. To interpret the importance of individual features in the classification, we used the SHAP feature importance method. Cellular intensity and vesicle numbers had a high average importance for classifying wortmannin and rapamycin treatments. Texture, intensity, and structure features of autophagosomes also contribute towards rapamycin classification. Additionally, we observed that rapamycin treatment initially leads to the formation of bigger autophagosomes. Our findings demonstrate that using SHAP to interpret machine learning models can assist in the identification of governing features and biologically relevant phenotypes from large datasets. While we used a model with satisfactory performance in identifying differentiating features between the three treatments, we acknowledge that the classifier accuracy could be further improved by optimizing hyperparameters or using other classification algorithms.

In this study, we focused on characterizing morphological phenotypes at a bulk level and did not fully leverage the benefits of the single cell temporal resolution. We observed heterogeneity in multiple features and time points for both rapamycin and wortmannin treatments. For example, wortmannin treatment caused a small portion of cells to have the opposite response compared to the rest for min_AP_contrast_mean and max_AP_area features at 30 minutes post treatment. Identifying features correlated to single cell heterogeneity could reveal novel insights. Coupling this kind of analysis with additional autophagy-related biosensors could also elucidate biological mechanisms involved in heterogeneity[25,26].

Image-based profile correlation between a standard drug (here rapamycin) and other treatments showed that temporal change in autophagy regulation can be effectively captured. For example, the correlation between rapamycin and DMSO profiles could be used as a threshold and any treatment with a higher correlation would represent an autophagy inducer or vice-versa. Similarly, for identifying autophagosome fusion modulators or autolysosome degradation modulators, additional reference drugs that affect those specific steps will be needed for comparison. However, it is worth noting that there could be pathway specific morphological variations between modulators that affect autophagy similarly. For instance, while rapamycin and another treatment may induce autophagy, they may not necessarily affect the same morphological features. Profiling morphological changes induced by a wider range of perturbations would aid in identification pathway-specific morphological alterations, as well as features that change globally in response to perturbations with a similar effect on autophagy. This could also reveal how distinct mechanisms lead to the same cellular output.

A simple profile similarity approach was used over more complex machine learning models because of the limited amount of data and to avoid overfitting. For instance, PI3K inhibitors and ULK1 inhibitors both inhibit autophagy, but might have different effects on the morphology. Therefore, a model trained on PI3K inhibitors may not accurately predict ULK1 inhibitors. In other words, machine learning models require richer training data and identification of features that are universally correlated with a general class of perturbations to achieve accurate prediction. Conversely, a profile similarity approach does not rely on pre-weighted features, which allows capturing the diverse differences in morphology. Nevertheless, machine learning models could be incredibly useful to predict autophagy states. Training a model at various concentrations could be used to predict how changes in autophagy impact more complex cellular decisions, such as cell death, at a single cell level.

A comparison of performance between pathway-specific and pathway-agnostic phenotypic characterization for drug screening would be an interesting avenue of analysis. The autophagy-related phenotypic characterization performed in this study is a pathway-specific phenotypic characterization approach, while assays such as Cell Painting are pathway agnostic[27]. Assays such as Cell Painting offer an unbiased and comprehensive measurement of cellular phenotypes. Such holistic measurements can capture any off-target effects that the drug might have, which can be overlooked if a specific pathway is monitored and targeted alone. Conversely, Cell Painting is limited by the temporal resolution it can achieve, which might be important for biological processes that are highly dynamic such as autophagy. Therefore, studies comparing the performance of both methods would be valuable in determining the optimal approach for characterizing perturbations.

In conclusion, image-based profiling of autophagy is an exciting approach that can be applied in various contexts to improve fundamental understanding of the autophagy pathway as well as expedite drug screening for various disease indications.

## Author Contributions

NSB conceived the work, executed experiments, developed image and data analysis pipelines, visualized data, wrote, and edited the manuscript. ET developed image analysis pipelines. PSS conceived the work, secured funding, wrote, and edited the manuscript.

## Acknowledgments

We thank Shiaki Arnett Minami and Matthew Kenaston from Shah lab for feedback on the manuscript.

## Declaration of interests

The authors declare no competing interests.

## Funding

This work was supported by the W.M. Keck Foundation.

## Methods

### RESOURCE AVAILABILITY

#### Lead contact

Further information and requests for resources and reagent should be directed to and will be fulfilled by the lead contact, Priya S Shah

#### Materials Availability

Cell lines generated are available from the lead contact upon request.

#### Data and Code availability

- Raw feature data (Table S1) and preprocessed data (Table S2) is available on https://data.mendeley.com/datasets/mftcnms5rh/1
- Raw image data is available upon request.
- All original code for feature extraction, preprocessing, and analysis are available on https://github.com/shahlab247/ATG_morphological_profiling.
- Any additional information required to reanalyze the data reported in this paper is available on request.

## METHOD DETAILS

### Cell culture, Chemical treatments, and Live cell imaging

Cell culture, generation of the A549-pHluroin-mKate2-LC3 reporter cell line, image acquisition for live cell imaging, and chemical treatments using rapamycin, wortmannin, and DMSO were previously discussed [11]. Three individual replicates were performed for collecting data for rapamycin and wortmannin experiments. The third replicate consisted of technical duplicates. Any disturbed images were removed from analysis through manual inspection.

### Image analysis, feature extraction and interpretation

Cell masks were generated using the Cellpose algorithm [28]. A custom-trained model was used for extracting cell masks [Available on GitHub]. Individual cells were tracked using the cell masks and bTrack algorithm [29]. Custom-optimized parameters were used as input for btrack [Available on GitHub]. Cell tracking efficiency was confirmed manually for various independent experiments. An example track series is available on GitHub. After the division of a cell, one of the daughter cells continues to have a parent ID and these cells were followed for the entire time course. The other daughter’s cell information is discarded. This leads to the loss of immense data and could be improved in the future.

Puncta mask for autophagosomes and autolysosomes are created using a spot detection tool in NIS elements software. Spot detection in different channels was previously elaborated [11]. The main changes to this protocol were 1) the Contrast value for detecting puncta in the TRITC channel was changed to 6. 2) The detected spots were further dilated using Grow Bright regions to Intensity operation to capture different sizes of puncta. 3) Puncta present in GFP and TRITC channels were detected using BOTH = TRITC HAVING (TRITC AND GFP) binary operation. 4) Only red puncta (autolysosomes) were detected using TRITC SUB BOTH. The general analysis file is available on GitHub. The accuracy of the masks was confirmed manually for cells under various treatment conditions. The total cell count detected using cell masks was also compared to cell count based on nucleus count Hoechst staining. The puncta masks are then subject to the watershed algorithm to differentiate conjoined adjacent puncta. The raw images, cell masks, and puncta masks of cells that are fully tracked are then used for extracting features.

Features were extracted for three biological categories-Cell as whole, Autophagosome (AP) and autolysosomes (AL). GFP channel images were used for extracting autophagosome and cellular features. For AL, TRITC channel images were used for extracting morphological features. GFP channel images have a better signal to noise ratio compared to TRTIC, hence used for extracting cellular features. Cell, autophagosome and autolysosome masks generated as described above were used for feature extraction. Regionprops from skimage.measure was used for extracting some of the features[30]. The mean and range of all 14 Haralick texture features calculated in all directions were included as features. 25 Zernike moments were calculated using 0.5*major_axis_length of the object as radius. Hararlick features and Zernike moments were estimated using Mahotas package [31]. After calculating features for each vesicle, descriptive statistics of each feature for all the vesicles in an individual cell were calculated. The descriptive statistics include mean, median (50%), lower quartile (25%), upper quartile (75%), maximum value(max), and minimum value(min). AP and AL features have their specific prefix before the feature and the prefix before AP/AL refers to the descriptive statistic of that feature. For example, mean_AP_area refers to the mean AP area of the all the AP in that specific cell. The list of 949 features extracted along with their respective categorization as biological entity and morphological type for each feature is available on GitHub.

### Data preprocessing and standardization

Features containing NAN values were removed. Each individual feature for each cell was centralized using the median value of DMSO-treated cells (median (X_DMSO_)) and divided by 1.2532 times the mean absolute deviation (mad (X_DMSO_)) of that feature from the respective plate (equation is shown below). 1.2532 times the mean absolute deviation (mad (X_DMSO_)) is approximately equal to the standard deviation [32]. This standardized data was referred to as modified Z score. This approach was used to account for the batch effects between experiments.

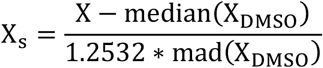

### Feature selection

All features with a median modified Z score ≥ 0.5 with an adjusted P-value < 0.05 were considered significant. Mann-Whitney U statistical test was used for estimating the statistical significance of each feature between DMSO-treated and small molecule treated cells. Benjamin-Hochberg method was used for false discovery rate correction.

### Flow cytometry

Cells were harvested and resuspended in PBS+ 1% FBS after treatment with DMSO and 100 nM rapamycin for the indicated time. The mean FITC intensity of the cells was analyzed using 488 nm laser on CytoFLEX S Flow cytometer. A minimum of 5,000 events were analyzed for each of the three independent replicates performed. Gating was applied to exclude 99.9% of non-fluorescent base strain A549 cells.

### Dimensionality reduction

2D UMAPs were generated using UMAP package on Python[33]. Hyperparameters used for generating the maps were neighbor=250 and mindist= 0.90.

### Random forest classification and feature importance

Features that varied significantly after treatment at all time points were combined at a single cell level. Features that were correlated with a Pearson correlation of 0.75 and above were removed except the first feature to remove redundancy. A lower correlation threshold of 0.75 was used to eliminate correlated features as co-linearity can considerably affect the feature importance interpretation [34]. RandomForestClassifier algorithm from sklrean.ensemble was used as the model. Random forest model accuracy was measured using a 5-fold cross-validation method. To elaborate, data was split into 5 folds, where 4 folds were used for training while the remaining fold was used for testing the accuracy of the prediction. We iteratively performed this step 5 times and calculated the average and standard deviation of the micro F1 score. 1000 trees were used as the random forest model, entropy as the criterion, and other hyperparameters are left as default. F1 micro score package from sklearn.metrics was used for calculating the F1 score [35]. A minimum of 70 cells were used for testing the accuracy of the model. Shapley Additive exPlanations (SHAP) method was used to interpret the feature importance from the classifier [18]. In short, SHAP assigns feature importance value for each feature by considering all possible combinations of features and the contribution of an individual feature to the final prediction. This was employed using SHAP package in Python.

### Hierarchical clustering and Pearson correlation

All features that varied significantly from DMSO-treated condition for all treatments at all time points were aggregated at a single cell level. In parallel, we used just the number of the autophagosome, and autolysosome features from all time points for comparison. Hierarchical clustering was performed based on median profiles of each treatment. clustermap function from seaborn package was used for clustering. ‘*average’* method and ‘*euclidean’* metric was used for generating the clusters. Pearson correlation was calculated using the median profiles of each condition.

## STATISTICAL ANALYSIS

Statistical tests used for determining significance are mentioned in the corresponding figure legend. The feature selection section describes statistical tests used for determining significantly variable features.

## Supplemental Information

**Figure S1:**
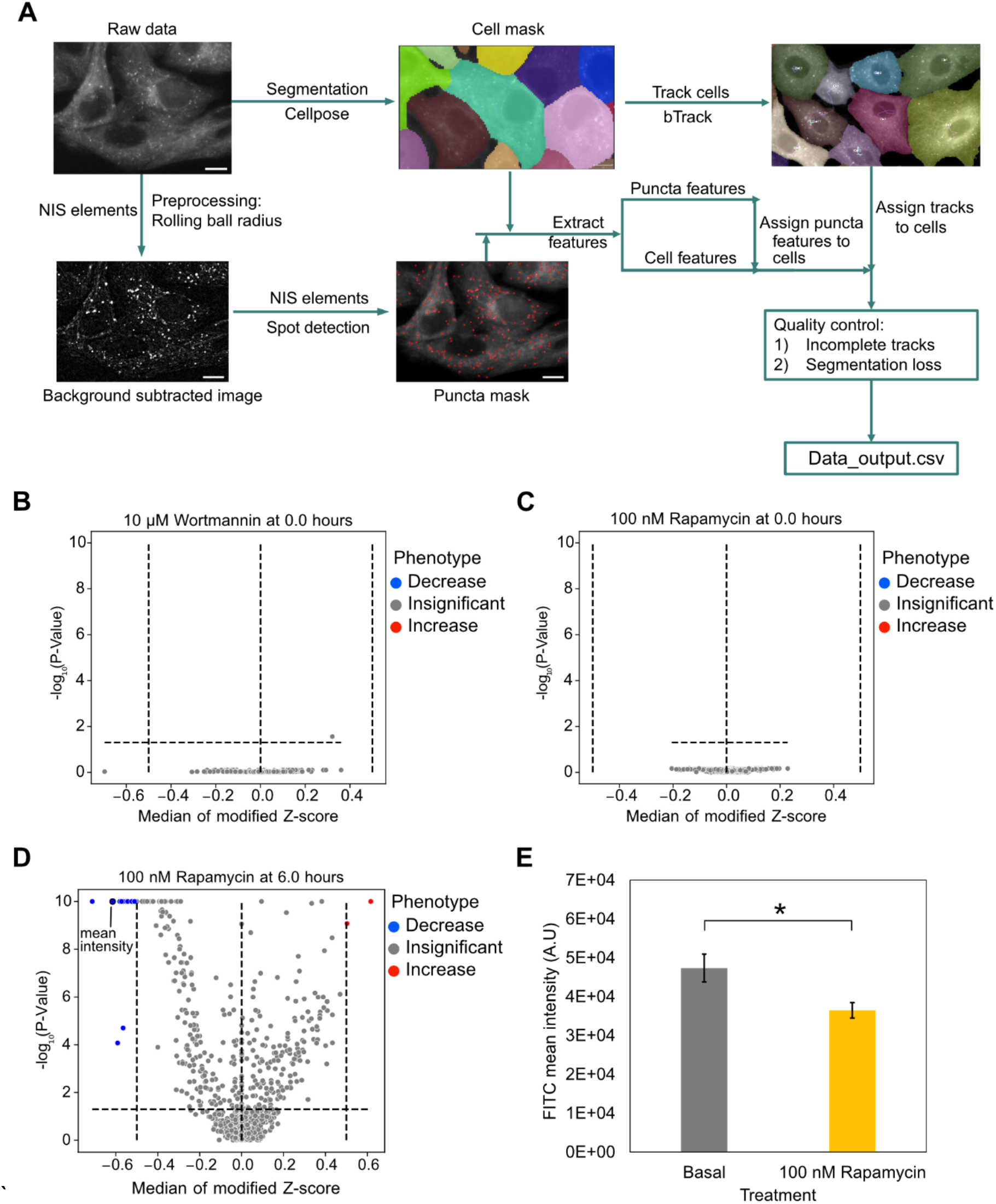
Image analysis pipeline and validation (related to Figure 1-2). **A**: Image analysis pipeline. **B-C:** Volcano plots of cellular features before treatment. **D:** Decrease in cellular mean intensity (GFP channel) measured using image analysis pipeline after treating with 100 nM rapamycin for 6 hours. **E:** Decrease in cellular mean intensity (FITC) measured using flow cytometry after treating with 100 nM rapamycin for 6 hours. Three independent replicates were performed. (^*^) represents p-value < 0.05. An independent two-tailed t-test was used to calculate the P-value.

**Figure S2:**
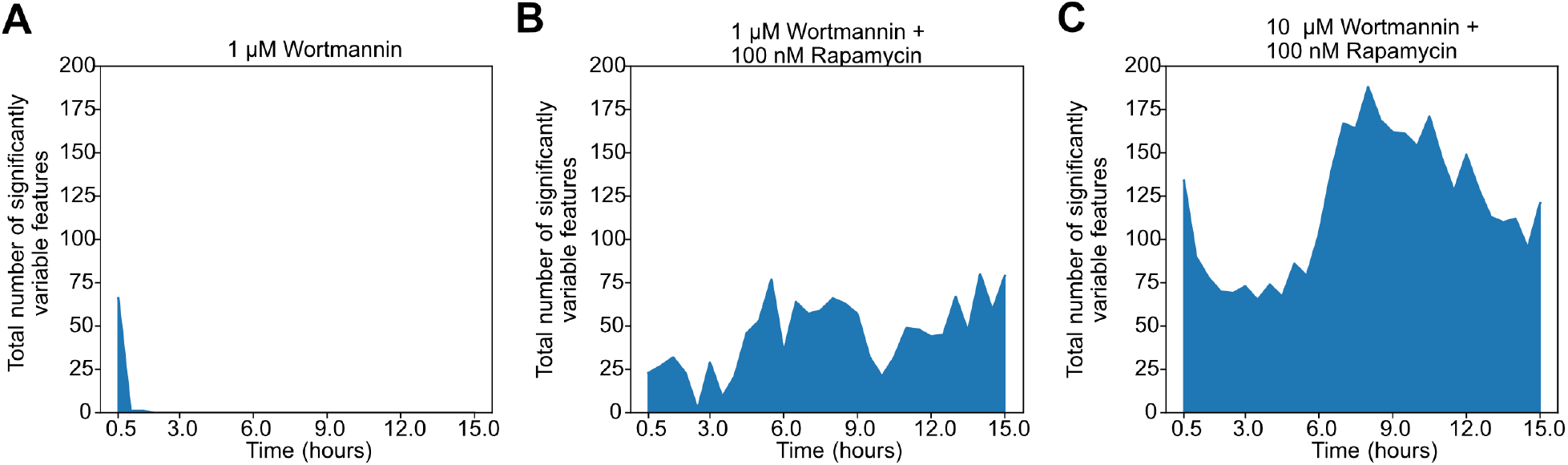
Concentration dependent effect of wortmannin on dynamic morphological features (related to Figure 2). Change in number of significantly variable features as a function of time after treating with **A:** 1 μM wortmannin **B:** 1 μM wortmannin and 100 nM rapamycin, and **C**: 10 μM wortmannin and 100 nM rapamycin.

**Figure S3:**
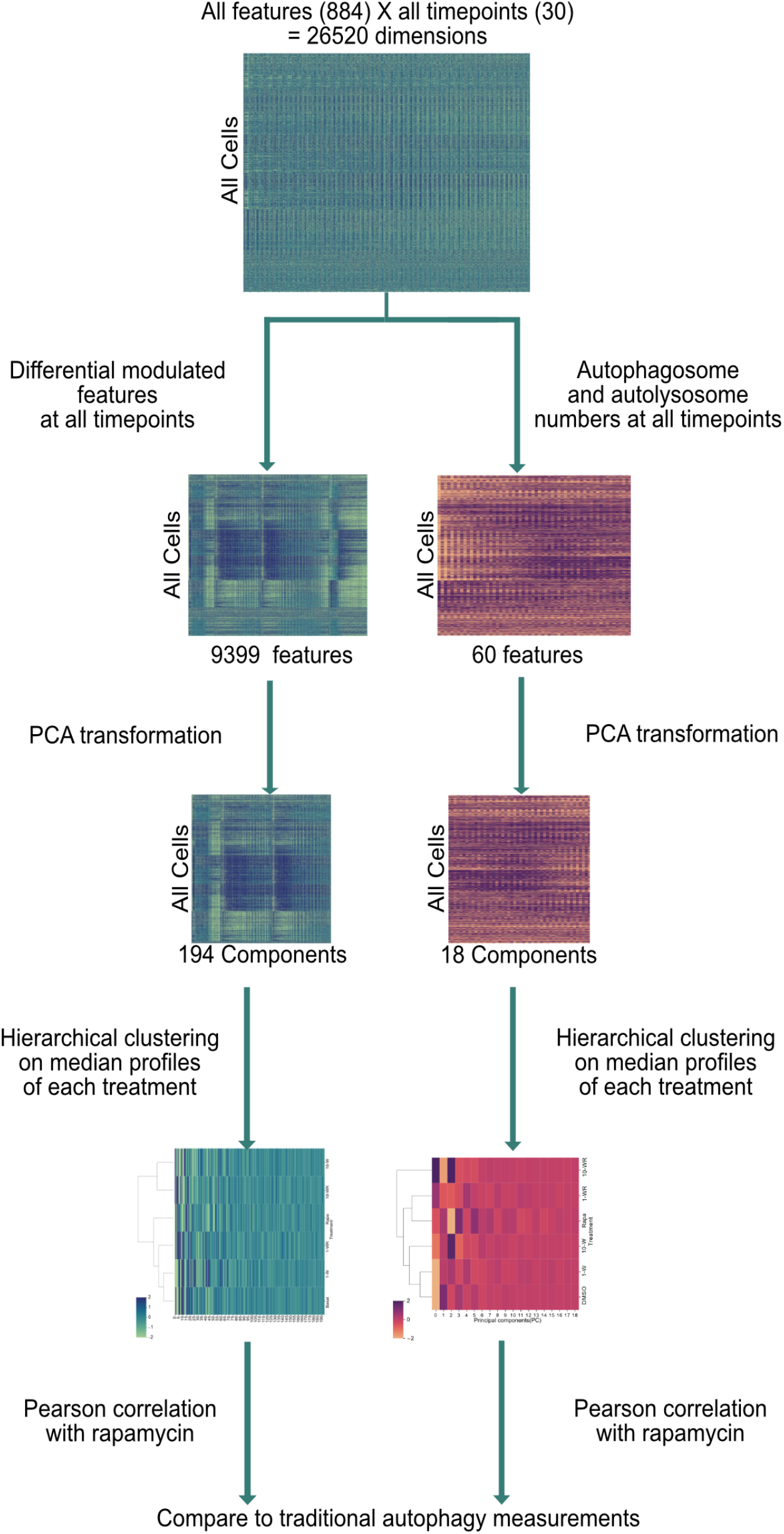
Stepwise procedure for calculating profile correlation (related to Figure 4). PCA was performed on features to reduce redundancy. 90% of the variance is retained after PCA. For hierarchical clustering based on median profiles, clustermap function from seaborn package was used. ‘*average’* method and ‘*euclidean’* metric was used for generating the clusters.

**Figure S4:**
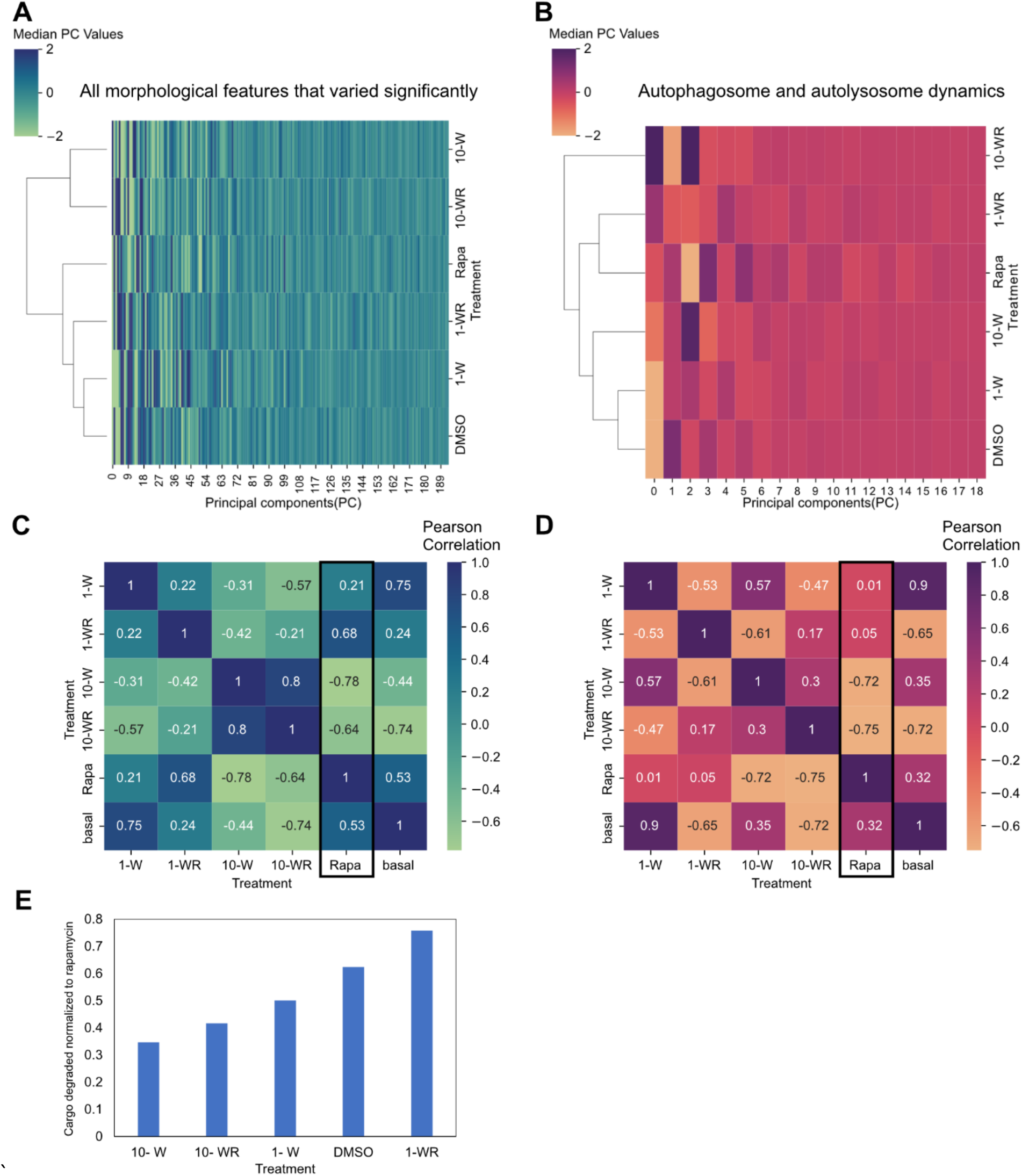
Assessing similarities among treatments using morphological profiles and autophagy dynamics (related to Figure 4). **A:** Hierarchical clustering of median profiles after principal components analysis (PCA) based on temporal image features. The color bar for heatmap represents median principal component (PC) values. **B:** Hierarchical clustering based on median profiles after PCA based on just temporal vesicle numbers. The color bar for heatmap represents median principal component (PC) values. **C:** Pearson correlation between profiles based on temporal image features. **D:** Pearson correlation between profiles based on just temporal vesicle numbers. **E**: Cargo degradation normalized to rapamycin treatment. Figure adapted from previous publication [1]. Abbreviations: 10 μM Wortmannin (10-W), 1 μM Wortmannin (1-W), 100 nM rapamycin (Rapa), 10 μM Wortmannin with rapamycin (10-WR), and 1 μM Wortmannin with rapamycin (1-WR).

## Notes

https://github.com/shahlab247/ATG_morphological_profiling

https://data.mendeley.com/datasets/mftcnms5rh/1

### Competing Interest Statement

The authors have declared no competing interest.

### Summary of Updates

Data availability is updated. Typos are corrected. Keywords are provided.

